# Increased nuclear factor I/B expression in prostate cancer correlates with AR expression

**DOI:** 10.1101/684472

**Authors:** Jagpreet S. Nanda, Wisam N. Awadallah, Sarah E. Kohrt, Petra Popovics, Justin M. M. Cates, Janni Mirosevich, Peter E. Clark, Giovanna A. Giannico, Magdalena M. Grabowska

**Affiliations:** Department of Urology, Case Western Reserve University, Cleveland, OH; Department of Pharmacology, Case Western Reserve University, Cleveland, OH; Department of Pathology, Microbiology, and Immunology, Vanderbilt University Medical Center, Nashville, TN; Department of Urology, Vanderbilt University Medical Center, Nashville, TN; Department of Urology, Levine Cancer Center/Atrium Health, Charlotte, NC; Department of Biochemistry, Case Western Reserve University, Cleveland, OH; Case Comprehensive Cancer Center, Case Western Reserve University, Cleveland, OH

**Author notes:** Address correspondence to: Magdalena M. Grabowska, 2123 Adelbert Road, Wood Research Tower; RTG00, Cleveland, OH 44106. Disclosure summary: The authors have nothing to disclose.

**Keywords:** prostate cancer, NFIB, androgen receptor

## Abstract

**Background:** Most prostate cancers express androgen receptor (AR), and our previous studies have focused on identifying transcription factors that modify AR function. We have shown that nuclear factor I/B (NFIB) regulates AR activity in androgen-dependent prostate cancer cells *in vitro*. However, the status of NFIB in prostate cancer was unknown.

**Methods:** We immunostained a tissue microarray including normal, hyperplastic, prostatic intraepithelial neoplasia, primary prostatic adenocarcinoma, and castration-resistant prostate cancer tissue samples for NFIB, AR, and synaptophysin, a marker of neuroendocrine differentiation. We interrogated publically available data sets in cBioPortal to correlate *NFIB* expression and AR and neuroendocrine prostate cancer (NEPCa) activity scores. We analyzed prostate cancer cell lines for NFIB expression via Western blotting and used nuclear and cytoplasmic fractionation to assess where NFIB is localized. We performed coimmunoprecipitation studies to determine if NFIB and AR interact.

**Results:** NFIB increased in the nucleus and cytoplasm of prostate cancer samples versus matched normal controls, independent of Gleason score. Similarly, cytoplasmic AR and synaptophysin increased in primary prostate cancer. We observed strong NFIB staining in primary small cell prostate cancer. The ratio of cytoplasmic-to-nuclear NFIB staining was predictive of earlier biochemical recurrence in prostate cancer, once adjusted for tumor margin status. Cytoplasmic AR was an independent predictor of biochemical recurrence. There was no statistically significant difference between NFIB and synaptophysin expression in primary and castration-resistant prostate cancer, but cytoplasmic AR expression was increased in castrationresistant samples. In primary prostate cancer, nuclear NFIB expression correlated with cytoplasmic NFIB and nuclear AR, while cytoplasmic NFIB correlated with synaptophysin, and nuclear and cytoplasmic AR. In castration-resistant prostate cancer samples, *NFIB* expression correlated positively with an AR activity score, and negatively with the NEPCa score. In prostate cancer cell lines, NFIB exists in several isoforms. We observed NFIB predominantly in the nuclear fraction of prostate cancer cells with increased cytoplasmic expression seen in castration-resistant cell lines. We observed an interaction between AR and NFIB through coimmunoprecipitation experiments.

**Conclusion:** We have described the expression pattern of NFIB in primary and castrationresistant prostate cancer and its positive correlation with AR. We have also demonstrated AR interacts with NFIB.

## Introduction

Since most prostate cancers express androgen receptor (AR), even advanced prostate cancer can be treated with androgen deprivation therapy^1^. Although these patients initially respond to androgen deprivation therapy, disease progression typically occurs within 30 months as these patients develop castration-resistant prostate cancer^2^. Castration-resistant prostate cancer is treated with newer androgen deprivation therapies such as abiraterone acetate (a steroidal CYP17A1 inhibitor^3^) and enzalutamide (a non-steroidal antiandrogen^4^)^5,6^. However, patients eventually progress on these second generation therapies as well and develop various castration-resistant phenotypes.

Approximately 63% of castration-resistant prostate cancers maintain AR protein expression^7^. In these tumors, AR signaling continues through multiple mechanisms including AR over-expression, increased androgen production, and induction of constitutively active AR splice variants (such as AR-V7)^8^. Prostate tumors can also escape androgen deprivation therapy by bypassing AR signaling. One mechanism is neuroendocrine differentiation, whereby cells acquire neuronal markers such synaptophysin and chromogranin A and sometimes lose AR expression and activity. Focal neuroendocrine differentiation is observed in 85% of castrationresistant prostate cancer patients^9^ while 8-25%^7,10–12^ of heavily treated prostate cancer patients, including those treated with abiraterone or enzalutamide, will undergo neuroendocrine differentiation to develop therapy-induced neuroendocrine prostate cancer^11,13–19^, which arises from the transdifferentiation of the adenocarcinoma^10,15,18,20–23^. Foci of neuroendocrine differentiation and therapy-induced neuroendocrine prostate cancers can be AR negative, have reduced AR activity, and be less responsive to androgen deprivation therapy^11,24^. In mouse models, neuroendocrine tumors can secrete neuronal factors, driving ligand-independent AR activation^25^. Another mechanism of androgen-independence in castration-resistant prostate cancer cells is activation of the fibroblast growth factor (FGF) pathway^7^. These “double negative” tumors lack both AR and neuronal markers and represent 21% of enzalutamide and abiraterone resistant tumors^7^.

Our previous studies have focused on identifying AR co-factors, based on the premise that these co-factors might play critical roles in prostate cancer progression. Building on the studies of Gao *et al*, which determined Forkhead box A1 (FOXA1) interacts with AR to drive prostate-specific gene expression^26^ and that Foxa1 is required for proper murine prostate gland development^27^ and maintenance^28^, we identified NFI transcription factors (NFIA, B, C, X) as FOXA1-interacting proteins that modulate AR-target gene expression^29^. Previous studies had demonstrated a role for NFI family members in regulation of AR-responsive promoters^30,31^, AR-target gene expression in castration-resistant prostate cancer cells^32^, and that NFI-consensus sequences were associated with AR binding sites in prostate cancer cell lines and tissues^33^. Our subsequent transient knockdown studies in androgen-dependent LNCaP cells demonstrated that NFIB inhibits AR-target gene expression and that NFIB is frequently associated with AR binding sites^34^. However, little was known about the expression pattern of NFIB in prostate cancer. We therefore performed a tissue microarray study to define the expression of NFIB in prostate cancer and determine whether its expression was associated with time to biochemical recurrence, neuroendocrine differentiation, or castration-resistant disease.

## Materials and Methods

### Human prostate cancer samples

Following Institutional Review Board (IRB) approval (Vanderbilt: 140838; CRWU: 20180111), we analyzed a prostate cancer tissue microarray linked with clinical data derived from the Urologic Outcomes Database at Vanderbilt University Medical Center. This tissue microarray has been previously described^35^. Our cohort included evaluable samples from 56 primary prostate cancer patients who had undergone radical prostatectomy and 8 castration-resistant prostate cancer patients who underwent tumor debulking procedures. The tissue microarray included replicate 1 mm cores from prostatectomy specimens and included normal prostatic tissue, benign prostatic hyperplasia (BPH), high-grade prostatic intraepithelial neoplasia (HGPIN), and adenocarcinoma, Gleason patterns 3-5. The tissue microarray was stained for NFIB (HPA003956, Sigma), AR (clone N-20, sc-816, Santa Cruz), synaptophysin (Clone 2, BD Biosciences) as a marker of neuroendocrine differentiation, and hematoxylin and eosin (H&E). Gleason score and immunohistochemistry staining was scored by a genitourinary pathologist (GG). H&E stained array cores were scored as normal, BPH, HGPIN, or Gleason pattern 3-5. Immunohistochemistry was scored by intensity (0 negative, 1 weak, 2 moderate, 3 strong) and distribution (0 negative, 1 <33%, 2 33-66%, and 3 >66%) in both nuclear and cytoplasmic compartments of luminal cells (normal, BPH) or tumor cells (HGPIN, prostate cancer, castration-resistant prostate cancer). A composite staining score was generated by summation of intensity and distribution scores (range 0-6). For correlation analysis and staining by Gleason score, we generated average values of duplicate cores. Unique cores and duplicate cores with differing Gleason scores were treated as unique cores. We also generated average per patient scores, averaging all cores to generate an average value for NFIB, AR, and synaptophysin across all conditions. These patient scores were used to evaluate expression across disease states and evaluate prostate cancer outcomes. Statistical analysis was performed using non-parametric tests, detailed in the figure legends. Following IRB approval (Vanderbilt: 160346; CRWU: 20180111), we also analyzed 8 de-identified patient samples of primary small cell prostate cancer obtained from O.U.R. Labs in 2005, with 1-3 needle biopsies available for analysis, as previously described^36^. Images were acquired using an Olympus BX43 microscope and a SC50 5 mega Pixel color camera and CellSens software. Immunohistochemistry images were white-balanced using Adobe Photoshop by using a tissue-less point on the slide as the reference.

### Hematoxylin and Eosin staining

Slides were soaked in xylenes (2 x 4 minutes), 100% ethanol (2 x 3 minutes), 95% ethanol (2 x 3 minutes), 70% ethanol (1 x 3 minutes), and 50% ethanol (1 x 3 minutes). Slides were then washed in running tap water for 3 minutes and dipped 10 times in de-ionized water. Slides were then placed in Harris hematoxylin (Richard Allan Scientific) for 4 minutes, followed by 3 minutes in running tap water and 10 dips in de-ionized water. Slides were then placed in Clarifier-2 (Richard Allan Scientific) for 45 seconds and rinsed 3 minutes in running tap water and 10 dips in de-ionized water. Rinsed slides were then placed in Bluing Reagent (Richard Allan Scientific) for 30 seconds and again rinsed 3 minutes in running tap water and 10 dips in de-ionized water. Slides were then placed in eosin-phloxine working solution (780ml 95% ethanol, 40ml 1% eosin Y in distilled water, 10ml 1% phloxine B in distilled water, and 4ml glacial acetic acid) for 2 minutes, and washed through 5 100% ethanol baths, 10 dips each before finally being dehydrated in xylenes (3 x 4 minutes). Slides were cover-slipped with Cytoseal 60.

### Immunohistochemistry

All sections were processed the same day, with developing times set by positive controls. Tissue microarray slides were soaked in xylenes (2 x 5 minutes), 100% ethanol (2 x 2 minutes), 95% ethanol (2 x 2 minutes), 70% ethanol (1 x 2 minutes), 50% ethanol (1 x 2 minutes). Slides were then washed in running tap water for 3 minutes and placed in a container with citric acid-based antigen unmasking solution (H-3300, Vector). Slides were then placed in a pressure cooker and cooked on high pressure for 25 minutes. Following pressure release, slides were allowed to cool for 25 minutes prior to subsequent steps. Slides were washed in 1 X PBS (3 x 10 minutes). Endogenous peroxidase activity was blocked with a 20-minute incubation of slides in peroxidase blocking solution (2.5ml of 30% hydrogen peroxide into 250ml methanol). Slides were again washed in 1X PBS (3 x 10 minutes). The tissue microarray was outlined with a PAP pen, and blocking solution (50μl of horse serum in 3 ml PBS) was added to slides for 30-minutes at room temperature. Blocking solution was removed and primary antibodies (1:1000) were added to each slide and incubated at 4°C overnight. In the morning, slides were washed in 1X PBS (3 x 10 minutes) before secondary antibody (1: 200, Vector) was added for an hour at room temperature. While slides were washed in 1X PBS (3 x 10 minutes), A and B components (2 drops A, 2 drops B into 5 ml PBS, VECTASTAIN Elite ABC HRP Kit) were combined to incubate for 30 minutes. AB solution was then added to slides and incubated at room temperature for 60 minutes, followed by washes in 1 X PBS (3 x 10 minutes). Slides were then developed with Liquid DAB+ Substrate Chromogen System (1 drop DAB+ per ml of buffer, DAKO), quenched with water and counterstained as follows. Positive controls (prostate for AR, NFIB, brain for synaptophysin) were used to set the exposure time, and all slides in an antibody series were incubated for the same amount of time in DAB. Slides were washed for 5 minutes in running tap water and then were placed in Harris hematoxylin (Richard Allan Scientific) for 30 seconds, followed by 1 minute in running tap water and 10 dips in de-ionized water. Slides were then dipped in Clarifier-2 for 3 times and rinsed 1 minute in running tap water and 10 dips in de-ionized water. Rinsed slides were then placed in Bluing Reagent (Richard Allan Scientific) for 10 dips and again rinsed 1 minute in running tap water and 10 dips in de-ionized water. Slides were then dipped 10 times in 70% ethanol and dehydrated through 5 100% ethanol baths, 10 dips each before being dehydrated in xylenes (3 x 4 minutes). Slides were cover-slipped with Cytoseal 60.

### Human clinical gene expression data

To correlate *NFIB* expression with AR activity in metastatic castration-resistant prostate cancer, we used cBioPortal^37,38^ to interrogate a set of publically available data^39^ with associated outcome, AR activity, NEPCa activity and gene expression data. We selected patients who were enzalutamide and abiraterone naïve for whom gene expression and scores were available (N= 106). The AR activity score is a measure of how active AR is in these tissue on the basis of a 27-gene signature derived from 24-hour treated R1881 treated LNCaP cells^40^. Similarly, the NEPCa score is a measure of whether tumors express genes more consistent with a therapy acquired neuroendocrine prostate cancer (NEPCa) as measured by a 70 gene signature of differentially expressed genes between castration-resistant prostate cancer adenocarcinomas versus therapy induced neuroendocrine prostate cancers^10^.

### Cell lines

LNCaP (clone FGC, CRL-2876), JEG-3 (HTB-36), 22RV1 (CRL-2505), PC-3 (CRL-1435), and DU145 (HTB-81) cells were purchased from American Type Culture Collection (ATCC). C4-2B^41^ cells were provided by Drs. Ruoxiang Wu and Leland Chung (Cedars-Sinai). BHPrS-1,, NHPrE-1, and BHPrE-1 cells were provided by Dr. Simon Hayward (NorthShore Research Institute)^42,43^. BHPrS-1 cells were maintained in RPMI supplemented with 5% FBS and NHPrE-1 and BHPrE-1 cells in F12/DMEM 1:1 medium containing 5% FBS, 1% Insulin-Transferrin-Selenium supplement, 10 ng/ml epidermal growth factor, 0.4% bovine pituitary extract and 1% antibiotic-antimycotic mix (all from Gibco). LNCaP, C4-2B, and 22RV1 were cultured in 10% premium fetal bovine serum (FBS, Atlanta Biologicals S11150) in RPMI 1640 + L-glutamine (Gibco). JEG-3 and DU145 cells were cultured in 10% FBS in MEM + Earle’s salts and L-Glutamine (Gibco). PC-3 cells were cultured in 10% FBS in F-12K Kaighn’s modification + L-Glutamine (Gibco). Cell culture experiments were performed in FBS-containing media.

### 3D culture of benign prostate cells

3D cultures of NHPrE-1 and BHPrE-1 cells were established in 8-well chamber slides by plating 2,000 cells/well in medium containing 2% (vol/vol) Matrigel on top of an undiluted layer of Matrigel. These cultures were grown for 9 days and cells were released from Matrigel using Corning Cell Recovery Solution to prepare cell lysates for Western blot analysis.

### Plasmids

pCMV6-Entry (vector) and pCMV6-NFIB-myc-DDK (RC231240) were purchased from Origene. The AR construct, p5HbHAR-A, used to generate AR-V7 and AR-V9^44^, was provided by Dr. Scott Dehm (University of Minnesota).

### Western blotting

Proteins from cell lines were lysed in ice-cold RIPA buffer (120 mM NaCl, 50 mM Tris (pH 8.0), 0.5% NP-40 and 1 mM EGTA) containing 1 mM phenylmethylsulphonyl fluoride, 1 mM sodium orthovanadate, 0.1 μM aprotinin and 10 μM leupeptin and kept on ice for one hour. Supernatants were collected by centrifugation at 13,750 rpm for 20 min at 4°C. Protein concentration was quantified and equal amounts of protein were added to 2X SDS loading buffer with 5% beta-mercaptoethanol. Nuclear and cytoplasmic extracts of LNCaP, C4-2B, 22RV1, JEG-3, DU-145 and PC3 cells were prepared using NE-PER Nuclear and extraction reagents (Thermoscientific, 78833). Equal amounts of cytoplasmic and nuclear protein extracts were resolved on NuPAGE 4-12% Bis-Tris Gels and transferred to nitrocellulose membranes. Membranes were blocked in 5% skimmed milk in TBS/0.1% Tween-20 (TBST) for 1 hour at room temperature followed by incubation overnight at 4°C with indicated antibodies NFIB (HPA003956, rabbit antibody, 1:1000 dilution, Sigma); AR (ab74272, rabbit antibody, 1:1000 dilution, Abcam); lamin B1 (ab133741, rabbit antibody, 1:2000, Abcam); GAPDH (AM4300, mouse antibody, 1:5000, Invitrogen) and α-tubulin (ab7291, mouse antibody, 1:5000, Abcam). GAPDH was used as cytoplasmic loading control whereas lamin B1 was nuclear loading control. Following secondary antibody incubation, proteins of interest were visualized using ECL detection reagent (GE Healthcare, RPN3243). Images were acquired using a Bio-Rad Chemi Doc. Molecular weights were calculated using Bio-Rad Image Lab. Average molecular weights of NFIB were determined from three independent experiments.

### Transient transfections

400,000 (JEG-3, LNCaP, 22RV1) or 300,000 (C4-2B) cells were plated in 6-well plates. After overnight recovery, cells were transfected with Silencer Select Pre-Designed siRNAs either as negative control (non-targeting, Silencer Select Negative Control No. 1 and 2 siRNA) or NFIB-targeting siRNA constructs (Thermofisher, s9494, s9496) using 30 pmol of siRNA in 9 μl of Lipofectamine RNAi Max. Cells were transfected for 72 hours, collected, and analyzed by Western blotting as above.

### Co-immunoprecipitation

2.5 x 10^6^ JEG-3 cells were seeded per plate in 10 cm dishes. After 10 hours, cells were transfected with 10 μg of empty, NFIB (variant 2), full length AR plasmid alone, or in combination using polyethylenimine (PEI) (Polysciences, Warrington, PA, USA) at a ratio of 3:1. Cells were washed with 1X PBS 48 hours after transfection and dislodged with a cell scraper. Cells were lysed in 500 μL of ice-cold RIPA buffer (120 mM NaCl, 50 mM Tris [pH 8.0], 0.5% NP-40 and 1 mM EGTA) containing 1 mM phenylmethylsulphonyl fluoride, 1 mM sodium orthovanadate, 0.1 μM aprotinin and 10 μM leupeptin and kept at 4°C on a rotospin for 1 hr. Supernatants were collected by centrifugation at 13,750 rpm for 20 min at 4°C. 80 μL of lysate was kept aside for protein quantification and analysis of input samples. The remainder of the lysate was used for preclearing by incubating them with 30 μL of equilibrated Pure Proteome TM Protein A Magnetic beads (LSKMAGA10) for 1 hour on ice. Then 4 μL of NFIB antibody from Sigma (HPA003956) was added per precleared sample for immunoprecipitation and kept on rotospin overnight at 4°C. The next day, 50 μL of equilibrated Protein A Magnetic beads were added to each sample and incubated for 6 hours before washing three times with RIPA buffer. Proteins were eluted with 60 μL of 2X SDS PAGE loading buffer with 5% betamercaptoethanol. Input and bead samples were heated at 70°C for 13 min and analyzed as above.

### Statistical analysis

Statistical analysis was performed using SPSS 24 (IBM) or GraphPad Prism and utilized non-parametric tests, with P values smaller than 0.05 considered significant.

## Results

Our previous study of a small cohort of patients demonstrated that NFIB was frequently lost in the luminal cells of patients with severe benign prostatic hyperplasia^34^. To determine whether this was also true in prostate cancer patients, we stained a tissue microarray and several core biopsies with an antibody against NFIB (Figure 1, Supplemental Figure 1A, B), which we had previously validated in *Nfib* knockout mouse tissues^34^. In order to determine whether changes in NFIB expression were associated with changes in AR or acquisition of neuroendocrine features, we also stained the tissue microarray for synaptophysin. Staining was then scored by a board-certified genitourinary pathologist (GG).

**Figure 1:**
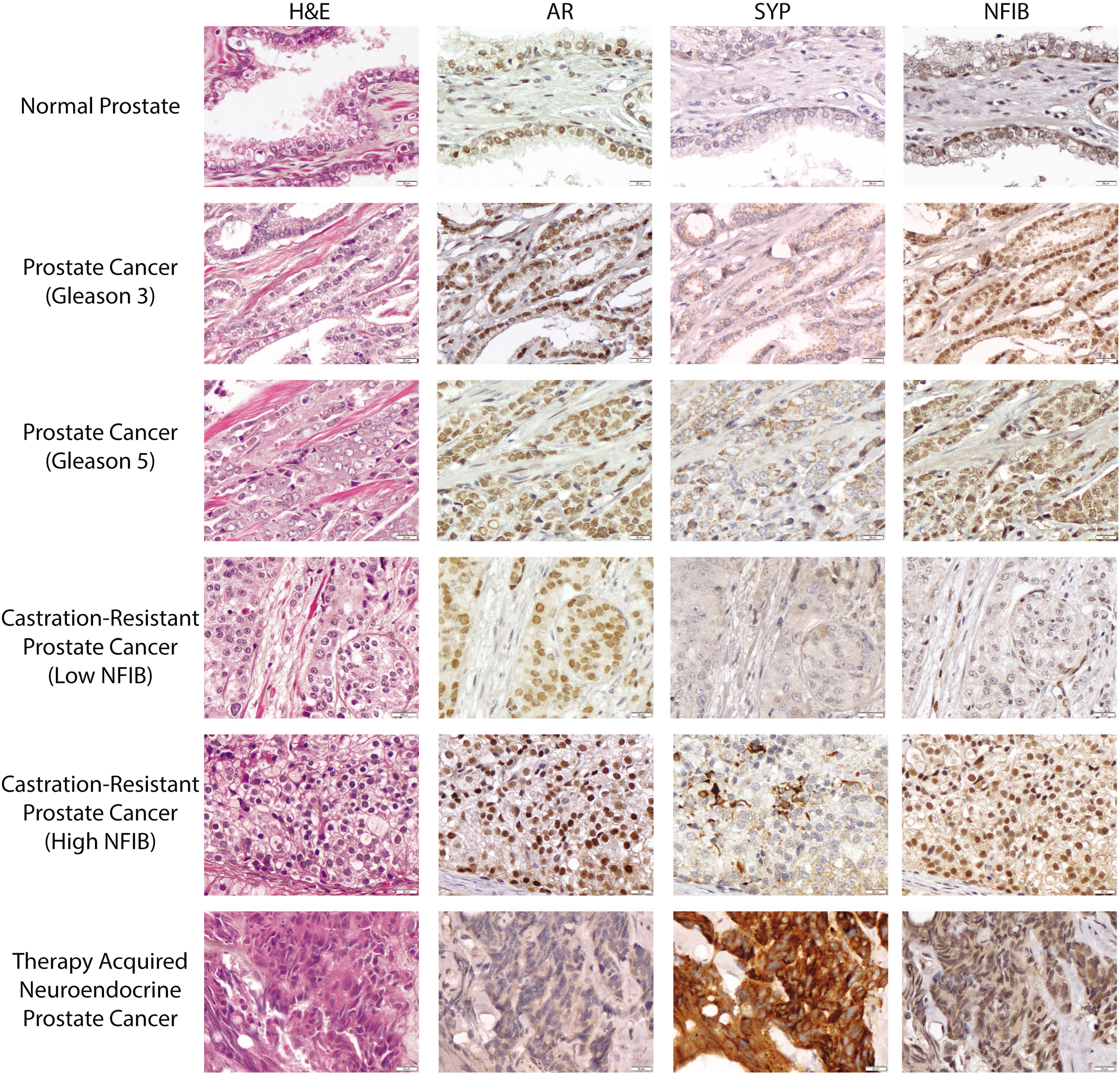
Expression of AR, synaptophysin, and NFIB in human prostate. Representative images from the human tissue microarray, representing normal prostate, prostate cancer, and castration-resistant prostate cancer. H&E: Hematoxylin and eosin; SYP: synaptophysin

Using an average patient score, we compared the expression of NFIB, AR, and synaptophysin in normal prostate and prostate cancer (Figure 2 A-C). NFIB was over-expressed in the nucleus (P = 0.0259) and cytoplasm (P < 0.0001) of cancer tissues versus patient matched normal prostate tissues. While AR was not over-expressed in the nucleus, cytoplasmic AR was increased in prostate cancer tissues versus normal prostate (P < 0.0001). Synaptophysin staining was limited to the cytoplasm and cell membrane, and also increased in prostate cancer tissues (P < 0.0001). To evaluate whether NFIB, AR, and synaptophysin expression corresponded with Gleason grade, we generated average scores per each unique core. There was no statistically significant difference between Gleason grade 3 versus 4 or Gleason grade 3 versus 5 for AR, NFIB, or synaptophysin (Supplemental Figure 2 A-C).

**Figure 2:**
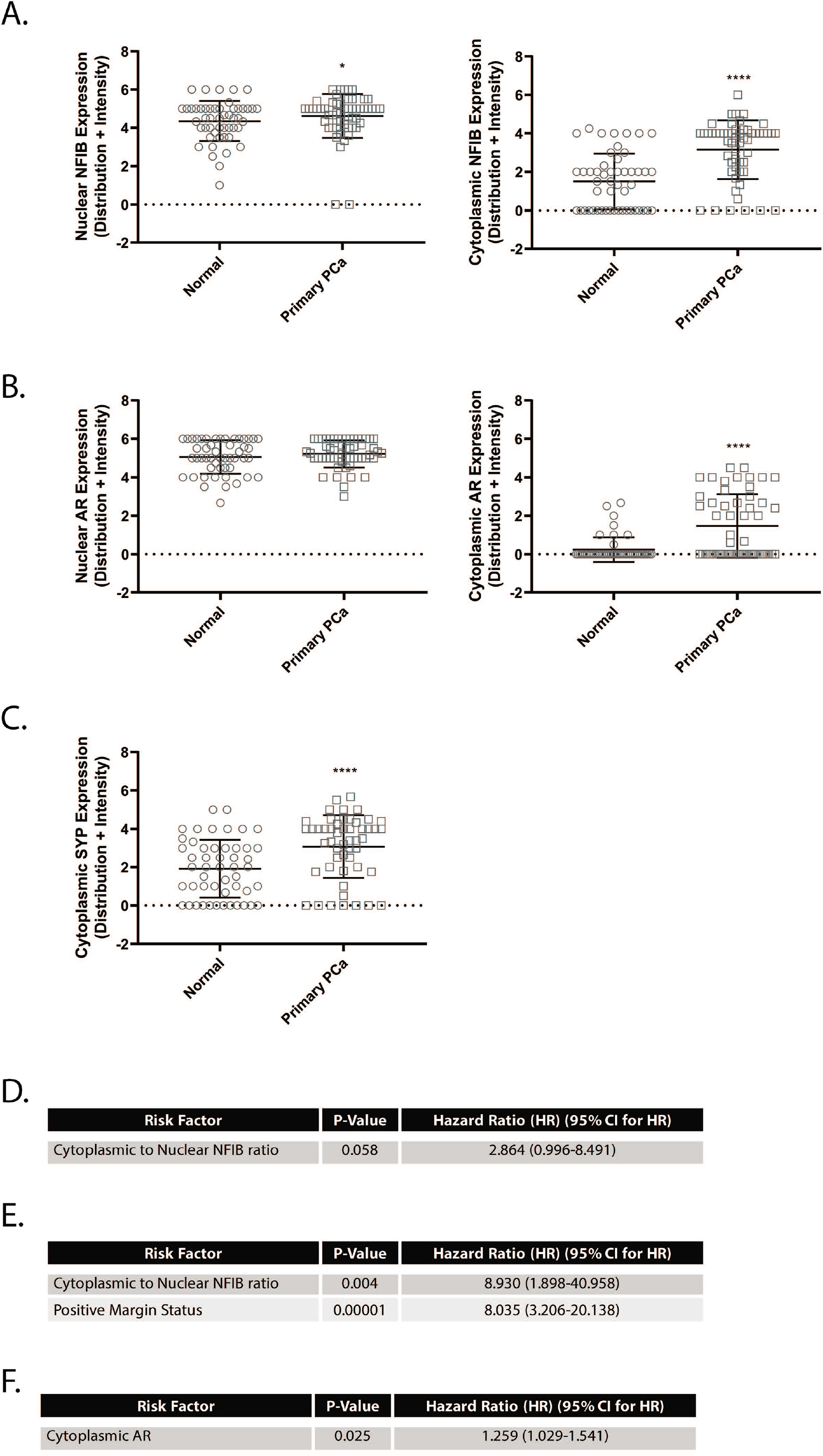
NFIB expression increases in prostate cancer and can predict earlier biochemical recurrence. Expression of NFIB (**A**), AR (**B**), and synaptophysin (**C**) in the nucleus and cytoplasm of normal prostate and prostate cancer. Intensity and distribution scores in the nuclear and cytoplasmic compartments of each core were added together to generate a composite expression score for each protein of interest and patient averages were generated (Matched patient N = 49 [NFIB], 47 [AR], 45 [SYP]). Data were analyzed using the Wilcoxon matched-pairs signed rank test. **D.** A high ratio of cytoplasmic-to-nuclear NFIB can predict earlier biochemical recurrence. Cox regression model predicting time to biochemical recurrence using the cytoplasmic-to-nuclear NFIB ratio. Analysis of 51 cases with 32 events (biochemical recurrence) and 19 censored cases. **E.** Cox regression model including positive margin status and cytoplasmic-to-nuclear NFIB ratio. Analysis of 49 cases with 31 events (biochemical recurrence) and 18 censored cases. **F.** Cytoplasmic AR can predict earlier biochemical recurrence. Cox regression model predicting time to biochemical recurrence using AR cytoplasmic staining. Analysis of 51 cases with 32 events (biochemical recurrence) and 19 censored cases. HR: hazard ratio CI: confidence interval; df: degrees of freedom. SYP: synaptophysin * P < 0.05; **** P< 0.0001

We also examined whether NFIB expression correlated with biochemical recurrence in 51 primary prostate cancer patients for who we had biochemical recurrence data. For these patients, we again used the average NFIB nuclear and cytoplasmic score. Nuclear and cytoplasmic NFIB staining did not correlate with biochemical recurrence (Supplemental 2D-F). However, when we examined a ratio of cytoplasmic and nuclear NFIB, we found a high ratio of cytoplasmic-to-nuclear NFIB predicted earlier biochemical recurrence (P = 0.058; hazard ratio 2.864; 95% CI 0.966 – 8.491; Figure 2D, Supplemental Figure 2G, H), an effect that achieved statistical significance when surgical resection margin status was accounted for in the Cox regression model (P = 0.004; hazard ratio 8.930; 95% CI 1.989 – 40.086; Figure 2E, Supplemental Figure 2H). There was no difference in cytoplasmic-to-nuclear NFIB by Gleason grade or tumor stage (Supplemental Figure–2J). Consistent with the literature^45^, we also observed that cytoplasmic AR staining could predict earlier biochemical recurrence (P = 0.025; hazard ratio 1.259; 95% CI 1.029-1.541; Figure 2F).

To more fully explore NFIB expression in the normal and diseased prostate, we compared NFIB average patient scores in unmatched patient samples, using unpaired analysis (Figure 3A, Supplemental Figure 3 A). Consistent with our previous report, NFIB was decreased in the nucleus of luminal cells in BPH-derived cores^34^, with low levels of cytoplasmic NFIB observed (Supplemental Figure 3A). While there was no statistically significant difference between nuclear NFIB staining in primary prostate cancer versus normal tissue or castration-resistant prostate cancer, cytoplasmic NFIB was again increased in primary prostate cancer versus normal prostate tissue (P < 0.0001) but not castration-resistant samples (Figure 3A). Notably, there appeared to be two types of castration-resistant prostate cancer patients: those with low nuclear NFIB and those with high nuclear NFIB expression (Figure 1, Figure 3A).

**Figure 3:**
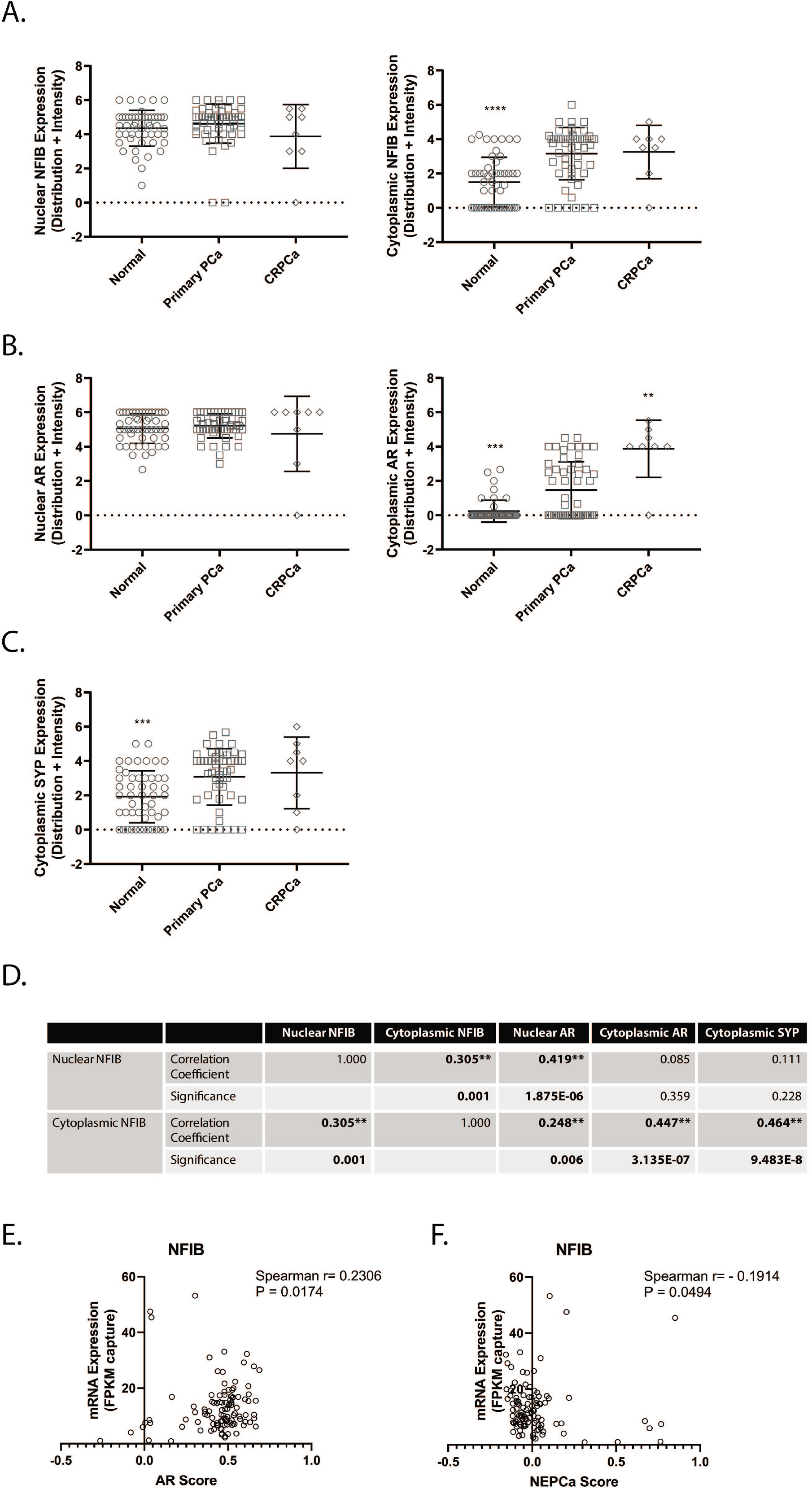
NFIB, AR, and synaptophysin expression and correlations in primary and castration-resistant prostate cancer. Expression of NFIB (**A**), AR (**B**), and synaptophysin (**C**) in the nucleus and cytoplasm of human prostate. Intensity and distribution scores in the nuclear and cytoplasmic compartments of each core were added together to generate a composite expression score for each protein of interest. An average value was generated for each patient. Significant values are reported between prostate cancer and normal prostate or castrationresistant prostate cancer. Data were analyzed using the Kruskal-Wallis test with Dunn’s multiple comparisons test. **D.** NFIB correlates with AR expression in primary prostate cancer. Intensity and distribution scores in the nuclear and cytoplasmic compartments of each core were added together to generate a composite expression score for each protein of interest and expression was correlated using Spearman correlation analysis (N= 120 unique cores). **E.** *NFIB* correlates with AR activity. Analysis of enzalutamide and abiraterone naïve (N= 106) castration-resistant prostate cancer patients samples from Abida *et al*^39^ analyzed in cBioPortal^37,38^ for *NFIB* and AR activity score via Spearman correlation analysis. **F.** *NFIB* inversely correlates with NEPCa activity. Analysis of enzalutamide and abiraterone naïve (N= 106) castration-resistant prostate cancer patients samples from Abida *et al*^39^ analyzed in cBioPortal^37,38^ for *NFIB* and NEPCa activity score via Spearman correlation analysis. PCa: prostate cancer; CRPCa: castrationresistant prostate cancer; SYP: synaptophysin ** P< 0.01; *** P< 0.001; **** P< 0.0001

We also examined AR and synaptophysin expression. While nuclear AR expression was unchanged, cytoplasmic AR was decreased in normal prostate samples (P = 0.0002) and increased in castration-resistant samples (P = 0.0038), versus primary prostate cancer (Figure 3B). Synaptophysin expression was also significantly lower in normal prostate tissue versus prostate cancer (P = 0.007), but there was no significant difference between primary and castration-resistant disease (Figure 3C).

Because only one of our castration-resistant prostate cancer cases satisfied our criteria for neuroendocrine features (loss of AR, strong synaptophysin staining), we also analyzed a limited number (n=8) of primary small cell prostate cancer core biopsies. Of these, 7 (88%) showed strong nuclear NFIB staining (Supplemental Figure 1B). Small cell prostate cancer cells have limited cytoplasm, so cytoplasmic NFIB expression was not scored.

To better understand the relationship between AR, NFIB, and synaptophysin in the more common prostatic adenocarcinomas, we also correlated the expression of nuclear and cytoplasmic NFIB with AR and synaptophysin. In primary prostate cancer samples, nuclear and cytoplasmic NFIB levels were positively correlated (Spearman rho = 0.305, P = 0.001), suggesting that NFIB expression increases in both compartments and that increasing cytoplasmic expression is not due to a shift in cytolocalization (Figure 3D). Nuclear NFIB was also positively correlated with nuclear AR (Spearman rho = 0.419, P < 0.0001). Cytoplasmic NFIB was positively correlated with nuclear AR (Spearman rho = 0.248, P = 0.006), cytoplasmic AR (Spearman rho = 0.447, P < 0.0001), and synaptophysin (Spearman rho = 0.464, P < 0.0001). The positive correlation of cytoplasmic NFIB with synaptophysin remained intact whether we examined all prostate cancer tissues (primary and castration-resistant) or castration-resistant only (Supplemental Figure 4A-E).

Since our immunohistochemistry data suggested there were NFIB-high and NFIB-low castration-resistant prostate cancer groups, and our castration-resistant prostate cancer sample size was rather limited, we turned to cBioPortal^37,38^ to interrogate a set of publically available gene expression data linked to AR and NEPCa activity scores^39^. We selected patients who were enzalutamide and abiraterone naïve (N= 106) and determined whether *NFIB* expression was associated with AR activity or NEPCa activity. Indeed, *NFIB* expression was positively correlated with AR activity (Spearman r = 0.2306, P = 0.017) and negatively correlated with NEPCa activity (Spearman r = −0.1914, P = 0.0494) (Figure 3E, F).

In order to evaluate whether established prostate cancer cell lines recapitulate the nuclear and cytoplasmic expression of NFIB, we performed nuclear and cytoplasmic protein isolation from prostate cancer cell lines. We examined androgen-dependent (LNCaP), castration-resistant (C4-2B, 22RV1), and AR-independent (DU-145, PC3) prostate cancer cells. As a control, we used JEG-3 cells which express limited NFIB based on previous reports and gene expression analysis^29,46^. NFIB isoforms were found in all cell lines (Figure 4A), including benign stromal and epithelial cells (Supplemental Figure 5A). For benign epithelial cells, there was no difference in NFIB isoform expression when grown in a monolayer or in 3D. Our antibody reliably identifies four bands in most prostate cancer cell lines, at 62 kDa, 57 kDa, 49 kDa, and 39 kDa (Supplemental Figure 5B), of which the 49 kDa band is the most prominent.

**Figure 4:**
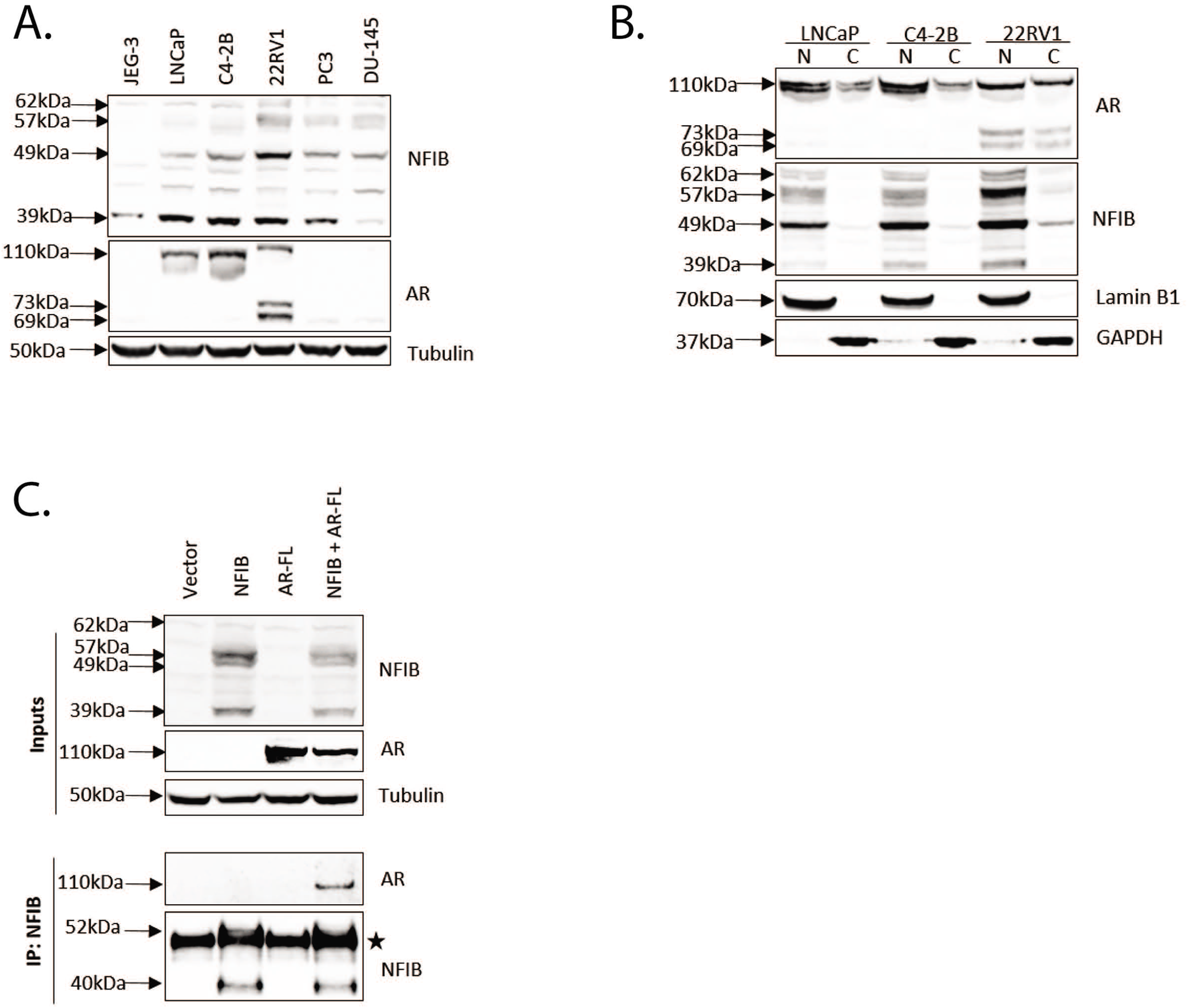
NFIB is expressed in prostate cancer cell lines and interacts with AR. **A.** NFIB expression in prostate cancer cell lines. Expression of NFIB in NFIB-low JEG-3 cells, androgen-dependent (LNCaP), castration-resistant (C4-2B, 22RV1), and androgen independent (PC3, DU-145) prostate cancer cells. **B.** Nuclear and cytoplasmic localization of NFIB. Nuclear and cytoplasmic preparations of LNCaP, C4-2B, and 22RV1 cells, analyzed for AR and NFIB expression. Nuclear loading control: Lamin B1; cytoplasmic loading control: GAPDH; N: nuclear; C: cytoplasmic **C.** NFIB interacts with AR. Transient transfection of JEG-3 cells with Vector, NFIB, full length AR (AR-FL), or AR-FL and NFIB, followed by co-immunoprecipitation with an antibody against NFIB and Western blotting for AR and NFIB. Star: antibody heavy chain IgG. IP: immunoprecipitation

In order to verify which of these bands are true NFIB bands, we performed transient transfections and over-expression studies. First, we transiently transfected PC-3 cells with NFIB over-expression or NFIB-knockdown constructs and compared which bands were impacted (Supplemental Figure 5C). In PC-3 cells, transient NFIB over-expression gave rise to a 53kDa band. Importantly, all other putative NFIB isoforms increased as well (62 kDa, 57 kDa, 49 kDa, and 39 kDa), suggesting that NFIB auto-regulates its own expression. Transient transfection of an NFIB-targeting siRNA also decreased expression of the 62kDa, 57 kDa, and 49 kDa bands, suggesting that these bands do indeed represent NFIB. We also expanded this analysis to JEG-3, LNCaP, C4-2B, and 22RV1 cells. In the LNCaP and C4-2B cells, transient knockdown abrogated expression of both the 57 kDa and 49 kDa bands (Supplemental Figure 5D). Neither of these were expressed in JEG-3 cells. In 22RV1 cells, NFIB knockdown also decreased expression of the 62 kDa band, which was not impacted in JEG-3, LNCaP, or C4-2B cells. However, as this band increased in response to NFIB over-expression and decreased in both 22RV1 and PC-3 cells in response to transient knockdown, we consider it an NFIB isoform. Finally, the 39 kDa band was not decreased in response to transient knockdown in the prostate cancer cells, but its expression was limited in JEG-3 cells and its expression was increased in response to transient over-expression of NFIB. We therefore cannot, at this time, determine whether it is an NFIB isoform.

Because cytoplasmic NFIB expression in prostate cancer patient tissues was higher in castration-resistant samples, we compared NFIB localization in LNCaP, C4-2B, and 22RV1 cells (Figure 4B). In LNCaP cells, which are androgen-dependent, NFIB is largely in the nucleus, but as cells progress to castration resistance in C4-2B cells, NFIB expression increases in both the nuclear and cytoplasmic fractions. These levels are even more pronounced in the unrelated castration-resistant prostate cancer cell line, 22RV1. Similarly, cytoplasmic AR expression increased in both castration-resistant prostate cancer tissue and cell lines.

Since AR and NFIB are frequently co-occupying the same cellular compartment, AR and NFIB co-occupy the same genomic regions in a prostate cancer cell line, and our previous studies determined NFIB regulates AR-target gene expression^29,34^, we evaluated whether NFIB and AR can interact directly by transfecting NFIB and AR into JEG-3 cells, which express low levels of NFIB. Indeed, in transient transfection co-immunoprecipitation experiments, NFIB and AR co-immunopurify (Figure 4C).

## Discussion

The goal of our study was to define the expression of NFIB in prostate cancer with respect to AR and synaptophysin and to begin to unravel the relationship between NFIB and prostate cancer progression. We analyzed a tissue microarray composed of cores including normal, BPH, prostate cancer, and castration-resistant prostate cancer tissues samples. On a matched per-patient basis, NFIB is over-expressed in both the nuclear and cytoplasmic compartments in primary prostate cancer versus normal prostate tissue. Importantly, increasing nuclear NFIB expression corresponds with increasing cytoplasmic expression, suggesting that NFIB continues to occupy the nucleus in prostate cancer. Indeed, there is no statistically significant difference between primary and castration-resistant prostate cancer in terms of nuclear or cytoplasmic NFIB expression.

Our observation of cytoplasmic NFIB is, to our knowledge, the first report of this transcription factor in the cytoplasm. However, another NFI family member, NFIC, has been reported in the cytoplasm of human dental cells where it localizes to the Golgi^47^ and helps regulate odontoblast differentiation^48^. The function on NFIB in the cytoplasm is presently unknown. It is possible that as it is over-expressed, it simply accumulates in the cytoplasm. This is supported by our correlation data, where increased cytoplasmic NFIB correlates with nuclear NFIB. Alternatively, cytoplasmic NFIB could be exerting non-genomic functions. In high-throughput screens, NFIB has been reported to interact with Dynactin Subunit 3 and SAS-6 Centriolar Assembly Protein^49^ and G protein-coupled receptor kinase 5 (GRK5)^50^. Notably, knockdown of GRK5 in androgen-independent prostate cancer cells resulted in decreased tumorigenicity^51^. Based on our interaction data, and the correlation of cytoplasmic AR and NFIB, it is possible NFIB is interacting with AR in the cytoplasm. In the cytoplasm, in response to androgens, AR can bind and activate Src, a non-receptor protein tyrosine kinase, resulting in activation of the Ras-Raf-MAPK/ERK cascade as well as activate the PI3K/AKT pathway (reviewed ^52,53^). What role NFIB plays in this process, if any, remains to be determined.

As there was not a significant difference between NFIB expression in either compartment when compared by Gleason pattern, we evaluated the utility of NFIB as predictor of biochemical recurrence, independent of Gleason grade. While nuclear NFIB or cytoplasmic NFIB expression alone could not predict biochemical recurrence, a higher ratio of cytoplasmic-to-nuclear NFIB predicted earlier time to biochemical recurrence (P = 0.058), and this achieved statistical significance when we accounted for surgical margin status in the Cox regression analysis. Surgical margin status is a complex variable as positive margins can arise from an imperfect surgery, more advanced disease, or combinations of both (reviewed^54^). Positive surgical margin status is an independent prognostic predictor of biochemical recurrence^54^, however the impact on metastasis and survival has been inconsistent, with some reporting negative predictive outcomes^55–57^ while others did not find any predictive value in surgical margin status in terms of survival and/or metastasis^58,59^. Notably, the extent of margin extension correlates with increased risk of biochemical recurrence^60–64^. In our analysis, we propose that adjusting for tumor margin status adjusts for more difficult cases/advanced disease, and supports the idea that a high ratio of cytoplasmic to nuclear NFIB is a predictor of biochemical recurrence following surgery.

This observation is exciting, but a limitation to our analysis is the sample size as well as the challenge of generating both a cytoplasmic and nuclear score for each patient. Moreover, the strongest predictor of biochemical recurrence in our study was positive margin status, which is evaluated after prostatectomy, limiting the utility of this biomarker at the time of biopsy. While these limitations limit the utility of NFIB as biomarker of biochemical recurrence, these observations do suggest NFIB plays an interesting role in prostate cancer biology.

Consistent with our human tissue data, NFIB is expressed in the nuclear and cytoplasmic fractions of prostate cancer cells. As in castration-resistant prostate cancer, castration-resistant prostate cancer cell lines have increased levels of NFIB in both the nuclear and cytoplasmic compartments compared to androgen-dependent prostate cancer cells. Consistent with UniProt predicted isoforms, we report that NFIB also exists in multiple isoforms in prostate cancer cells, with average molecular weights of 62 kDa, 57 kDa, and 49 kDa.

Our observation of multiple NFIB isoforms is novel in prostate cancer cell lines, but has been reported in other systems. The *NFIB* gene can undergo alternative splicing, giving rise to nine variants^65^, and the presence of multiple NFIB protein isoforms is consistent with reports from UniProt^66^, where NFIB has six reported isoforms (O00712 [NFIB_HUMAN], Entry version 172 [08 May 2019]). These isoforms have molecular weights of 47,442 Da (O00712-1), 22,251 Da (O00712-2), 53,049 Da (O00712-4), 55,181 Da (O00712-5), and 33,525 Da (O00712-6). While these do not match up exactly with our reported bands at 62 kDa, 57 kDa, and 49 kDa, NFIB a can undergo posttranslational modification like glycosylation^67^ and sumoylation^68^ which could add to these molecular weights. *In vitro* translation assays have also identified NFI-B2 and NFI-B3, at 47 kDa and 22 kDa, respectively^69^, indicating that some of these splice variants are expressed and functional. Significantly, the NFI-B3 isoform lacks the ability to bind DNA and regulate gene expression and acts as a dominant negative factor in the presence of NFIB, NFIC, and NFIX^69^. Unfortunately, the NFIB antibody immunogen is a sequence shared by all but the 22,251 Da (O00712-2) isoform. Therefore, we do not know the status of this isoform in our prostate cancer cell lines or prostate cancer tissues.

Although our previous studies showed that FOXA1 bridges the AR/NFIX interaction in HeLa cells in Fluorescent Protein Förster Resonance Energy Transfer (FP-FRET) experiments^29^, our exogenous co-immunoprecipitation studies now demonstrate at least one NFI, NFIB, is capable of interacting with the full length AR independently of FOXA1. What domains are responsible for this interaction, whether this interaction occurs with AR splice variants, and what the functional consequences of AR and NFIB interaction in primary and castration-resistant prostate cancer entail is under investigation.

Although our studies have focused on the role of NFIB in regulating AR expression and activity, it is likely that NFIB also has roles independent of AR. For example, in mouse models of small cell prostate cancer or lung cancer, *Nfib* is regularly amplified^70,71^, and over-expression of *Nfib* in a transgenic model of small cell lung cancer drives aggressive disease^72^. Our small cohort of primary small cell prostate cancer samples strongly express NFIB. What the role for NFIB in these AR-independent tumors is also remains to be determined.

In summary, our study has described the expression of NFIB, AR, and synaptophysin in a small cohort of prostate cancer patients. We report that NFIB is over-expressed in the nuclear and cytoplasmic fractions of prostate cancer patient tissues and cell lines. We also report that NFIB can interact with full length AR.

## Acknowledgements

We would like to thank the prostate cancer patients included in the tissue microarray for providing their tissues and clinical data. We would also like to thank Dr. Robert J. Matusik for his early support of these studies and critical feedback on the manuscript, and Daniel A. Barocas whose development of the Urologic Outcomes Database enabled linking outcomes data with this tissue microarray^35^. This work was supported in part by the Defense Health Program, through the Prostate Cancer Research Program Postdoctoral Training Award, under Award No. W81XWH-14-1-0312. Opinions, interpretations, conclusions and recommendations are those of the author and are not necessarily endorsed by the Department of Defense or U.S. Army. In the conduct of research utilizing recombinant DNA, the investigators adhered to NIH Guidelines for research involving recombinant DNA molecules. These studies were also supported in part by the National Cancer Institute of the National Institutes of Health under award number K99CA197315 and R00CA197315 (to MMG). The content is solely the responsibility of the authors and does not necessarily represent the official views of the National Institutes of Health.

---

**Supplemental Figure 1:**
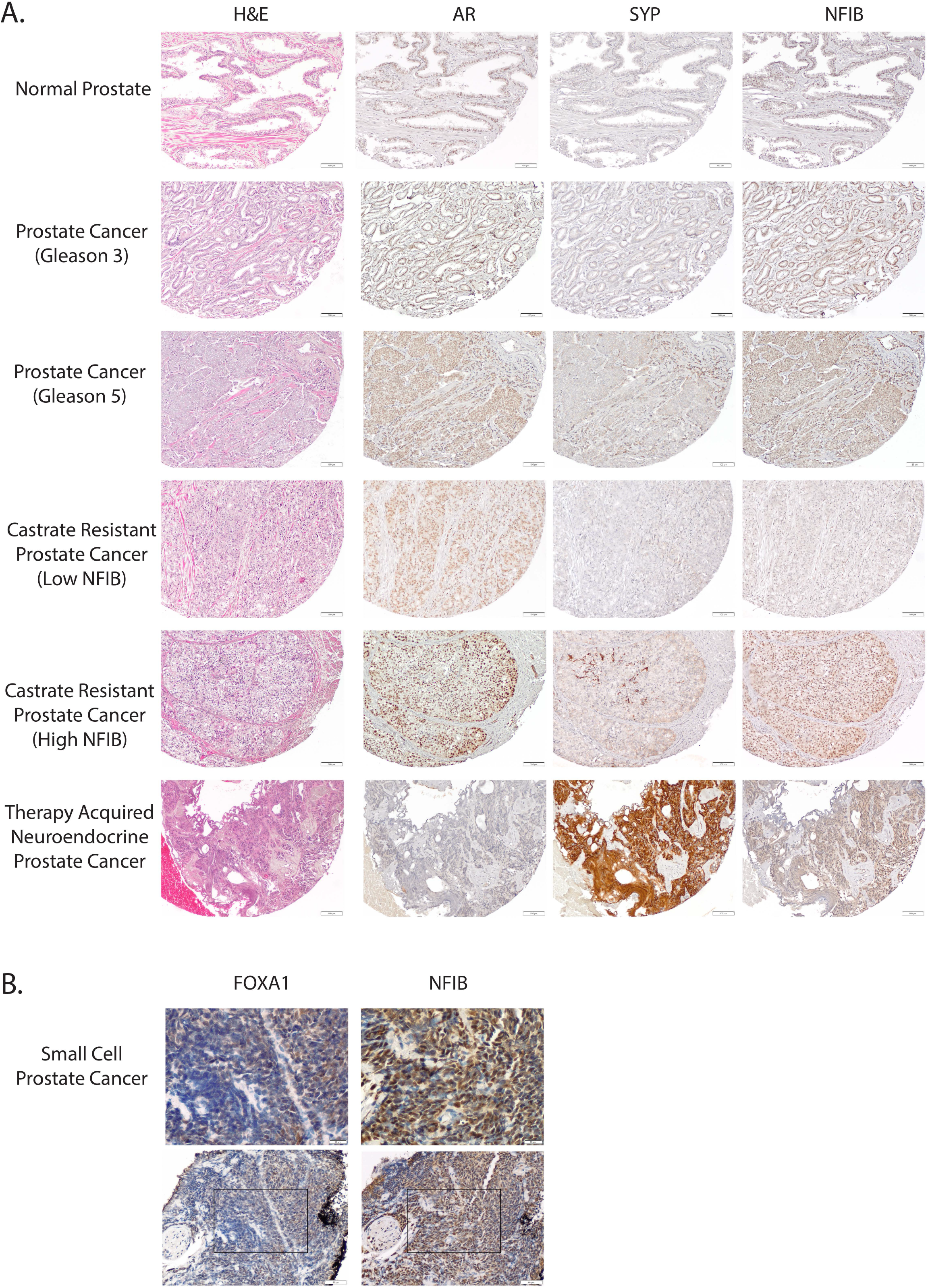
Expression of AR, synaptophysin, and NFIB in human prostate. **A.** Representative images from the human tissue microarray, representing normal prostate, prostate cancer, and castration-resistant prostate cancer. **B.** FOXA1 and NFIB staining in primary human small cell prostate cancer. Rectangle: area magnified in top panel. H&E: Hematoxylin and eosin; SYP: synaptophysin

**Supplemental Figure 2:**
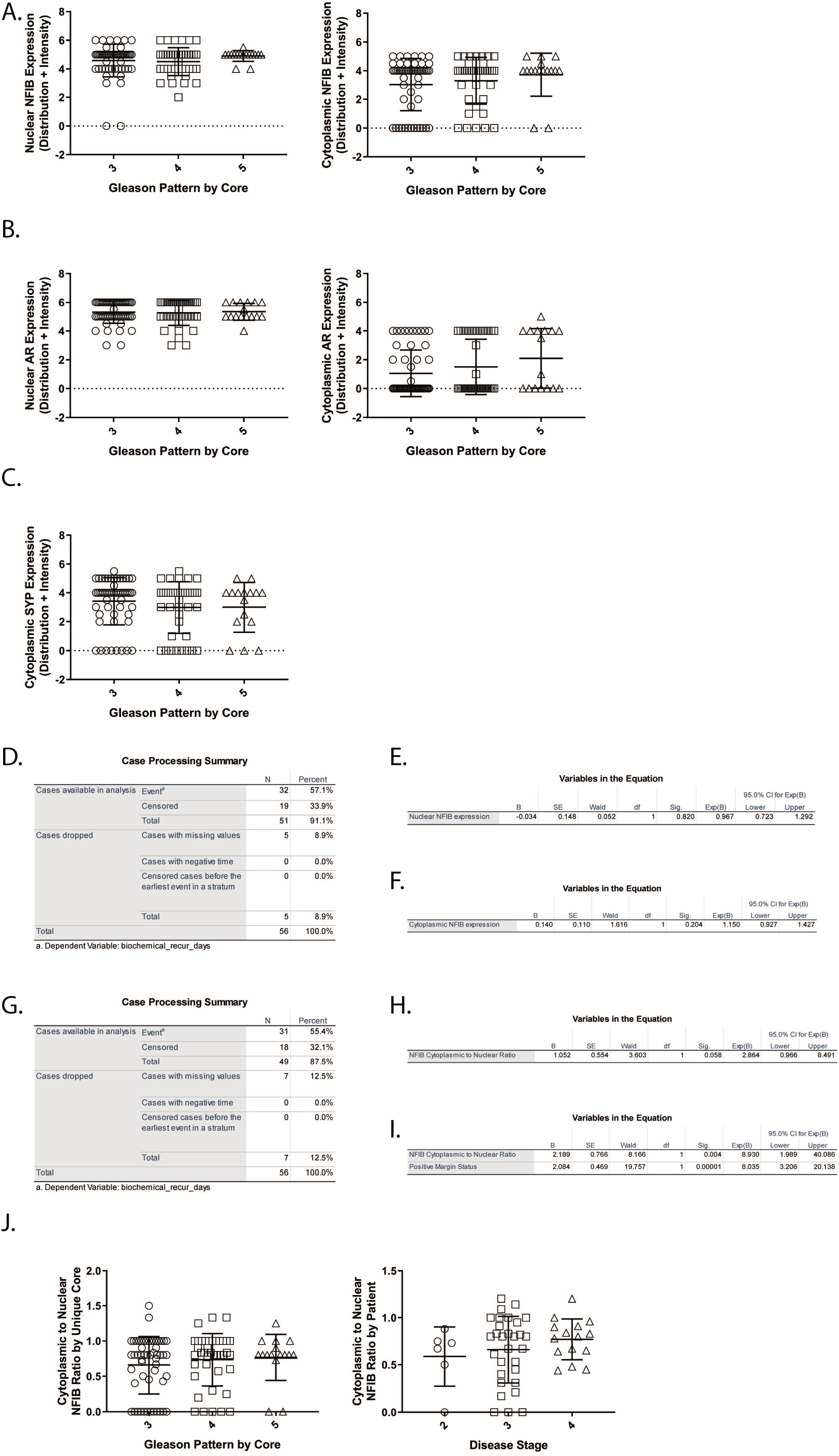
Expression of NFIB, AR, and synaptophysin across prostate diseases and prediction of biochemical recurrence. **A/B/C.** Nuclear and cytoplasmic AR, NFIB, and synaptophysin expression does not vary by disease grade or stage. For analysis of association of Gleason score with NFIB (**A**), AR (**B**), and synaptophysin (**C**) staining, unique core values were used (Unique core N = 56 Gleason 3, 40 Gleason 4, 16 Gleason 5). Significant values are reported relative to Gleason score 3. Data were analyzed using the Kruskal-Wallis test with Dunn’s multiple comparisons test. **D.** Case processing summary of NFIB as a single variable to predict time to biochemical recurrence (event) in days. **E/F.** Nuclear and cytoplasmic NFIB alone do not predict biochemical recurrence. Variables in the Cox regression model. **G**. Case processing summary of cytoplasmic-to-nuclear NFIB ratio and surgical margin status to predict time to biochemical recurrence in days. **H.** Cox regression model predicting time to biochemical recurrence using the cytoplasmic-to-nuclear NFIB ratio. **I.** Cox regression model including positive margin status and cytoplasmic-to-nuclear NFIB ratio. **J.** Cytoplasmic-to-nuclear ratio of NFIB does not vary by Gleason pattern by core or by patient tumor stage. B: unstandardized regression coefficient; SE: standard error; Exp (B): hazard ratio; CI: confidence interval; df: degrees of freedom, SYP: synaptophysin * P < 0.05; **** P< 0.0001

**Supplemental Figure 3:**
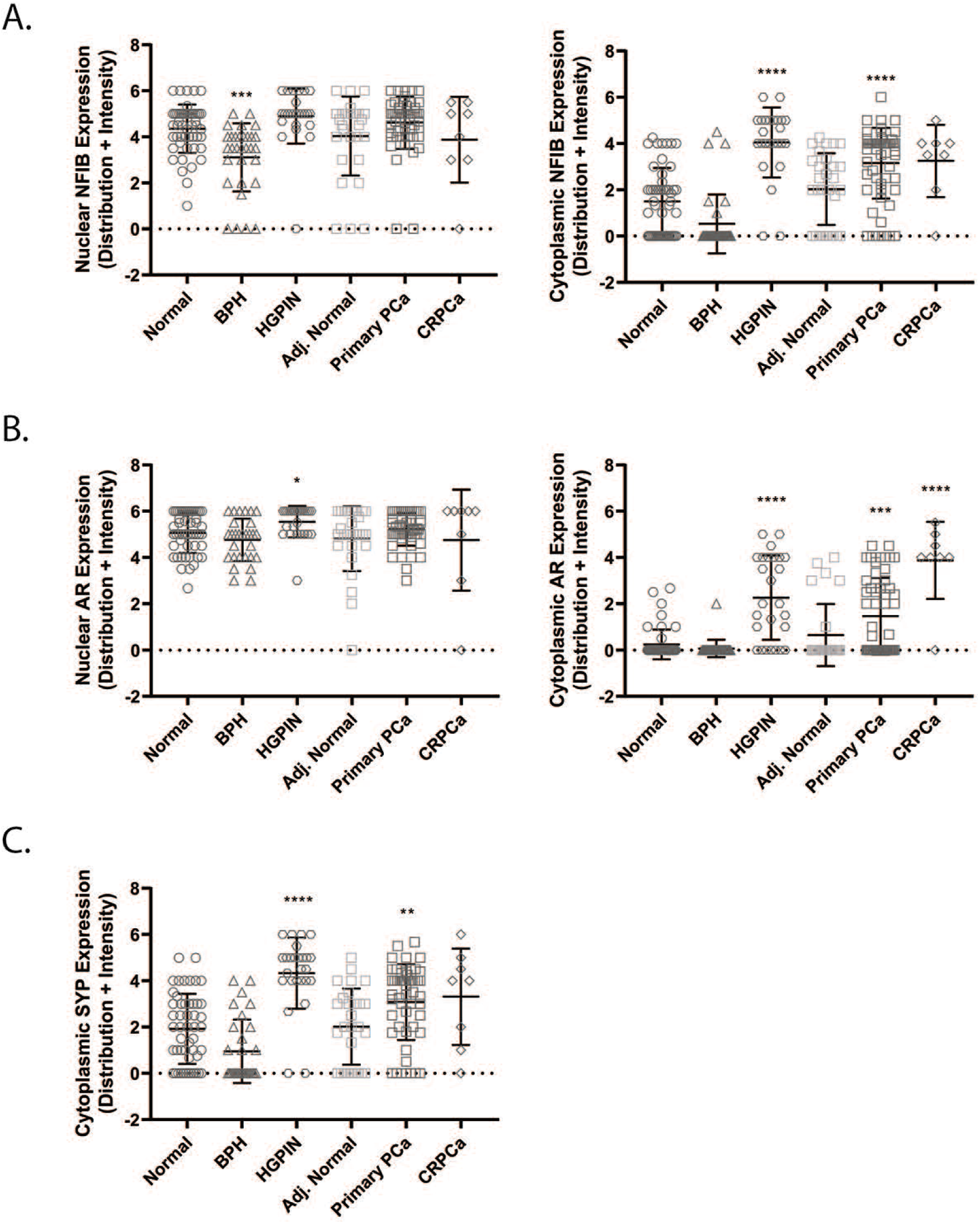
Expression of NFIB, AR, and synaptophysin in the prostate and in prostatic diseases. Expression of NFIB (**A**), AR (**B**), and synaptophysin (**C**) in the nuclear and cytoplasmic fractions of human prostatic tissues. Intensity and distribution scores in the nuclear and cytoplasmic compartments of each core were added together to generate a composite expression score for each protein of interest. Data were analyzed using the Kruskal-Wallis test with Dunn’s multiple comparisons test. Significant values are reported between normal prostate (normal) and diseased prostate. BPH: benign prostatic hyperplasia; HGPIN: high grade prostatic intraepithelial neoplasia; Adj. Normal: adjacent normal; PCa: prostate cancer; CRPCa: castration-resistant prostate cancer; SYP: synaptophysin * P < 0.05; ** P< 0.01; *** P< 0.001; **** P< 0.0001

**Supplemental Figure 4:**
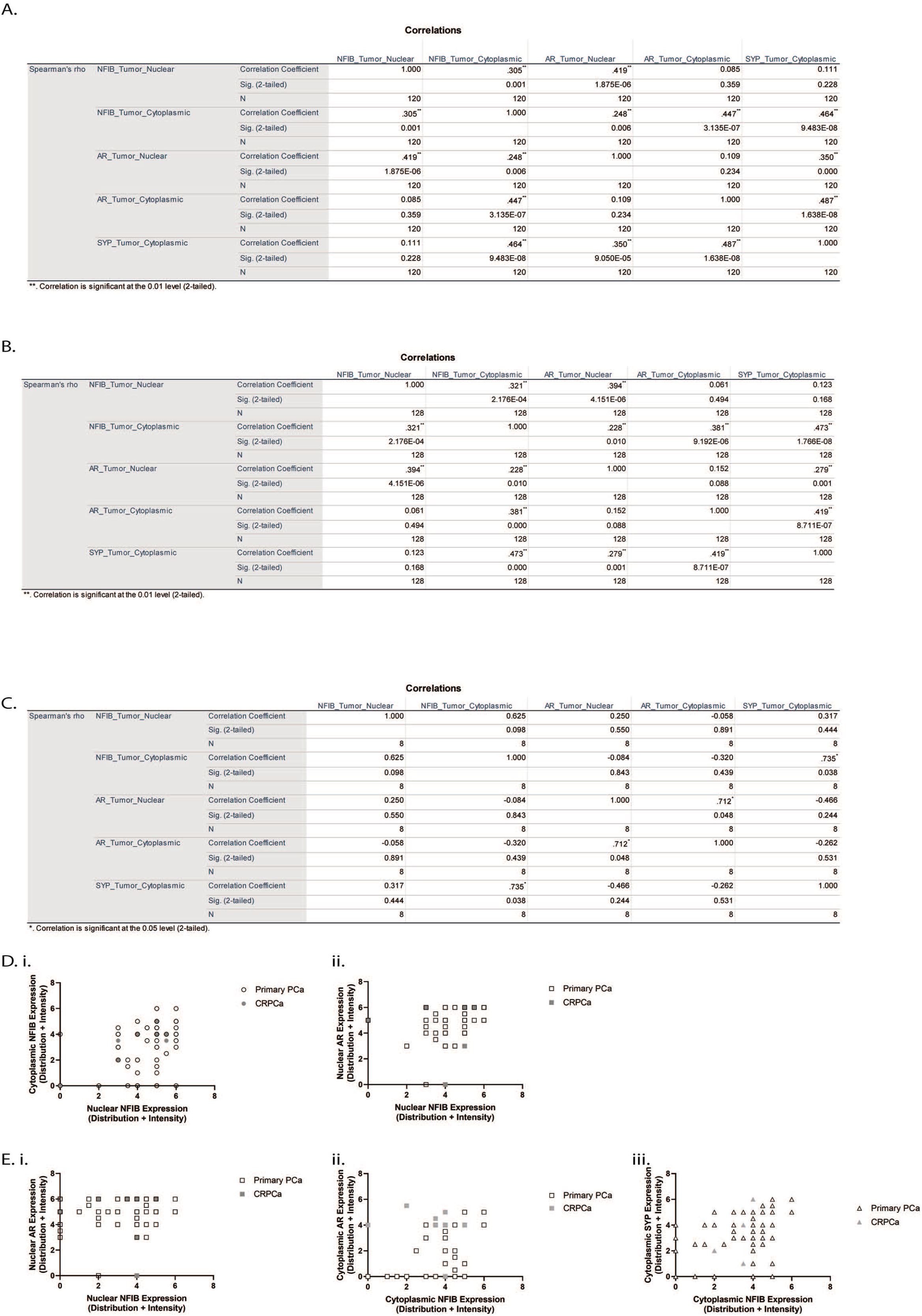
Correlations between NFIB, AR, and synaptophysin in prostate cancer. Spearman correlations between nuclear and cytoplasmic staining for NFIB, AR, and synaptophysin in primary prostate cancer (**A**), both primary and castration-resistant prostate cancer (**B**), and castration-resistant only (**C**). Graphical representations of significant correlations between nuclear NFIB (**D**) and cytoplasmic NFIB (**E**).

**Supplemental Figure 5:**
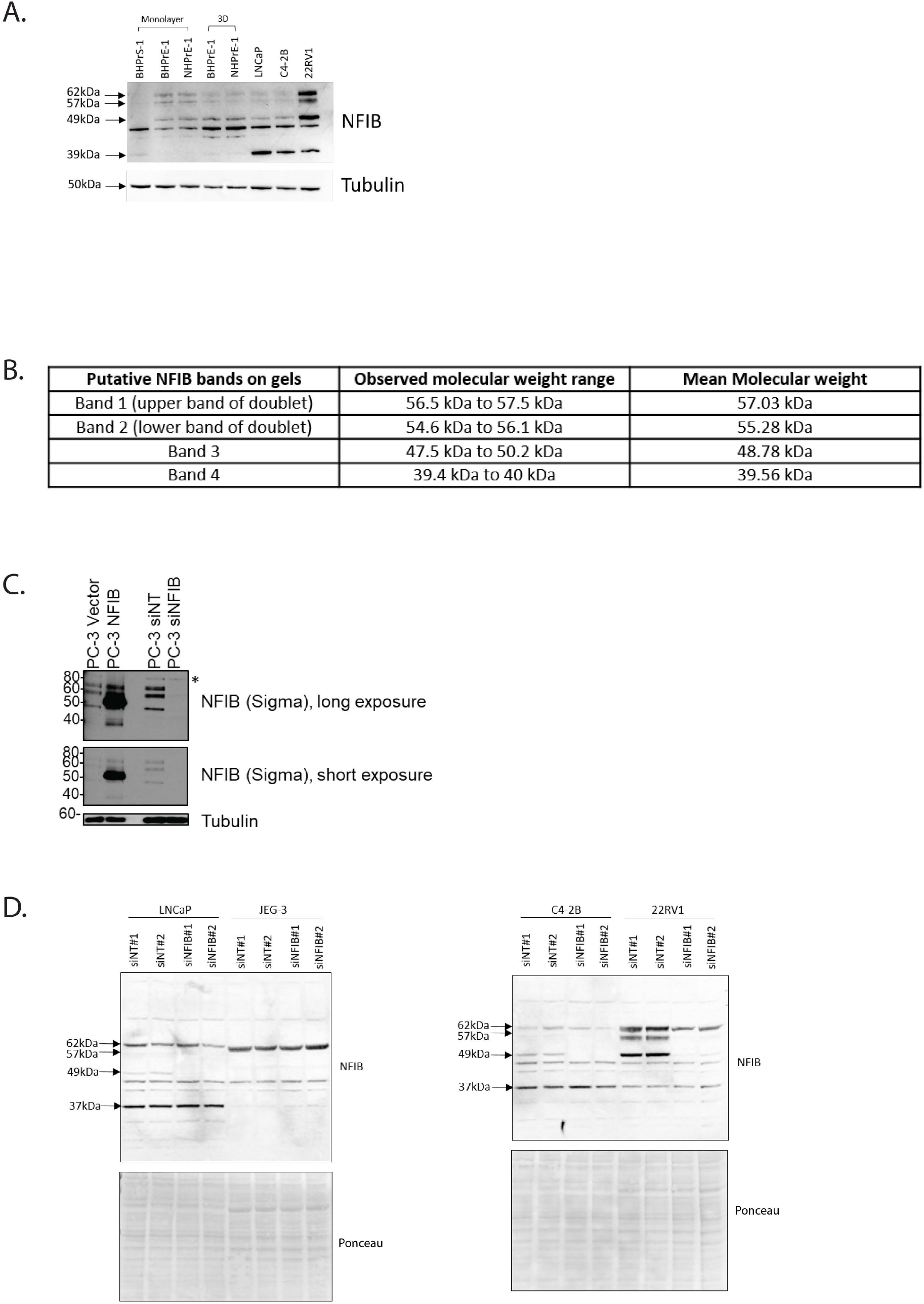
NFIB isoforms. **A.** Expression of NFIB in stromal and epithelial cells derived from BPH patients (BHPrS-1, BHPrE-1) and non-hyperplastic prostate epithelial cells (NHPrE-1). Cancer cell lines shown as control. **B.** Calculated molecular weights of NFIB. **C.** NFIB exists in several isoforms in PC-3 cells. Transient over-expression of vector or NFIB or knockdown of non-targeting control RNA or NFIB in PC-3 cells. **D.** NFIB exists in several isoforms. Transient knockdown of non-targeting siRNA or NFIB in JEG-3, LNCaP, C4-2B, and 22RV1 cells. Ponceau S staining for total protein serves as a loading control.

